# Mean Shift Clustering as a Loss Function for Accurate and Segmentation-aware Localization of Macromolecules in Cryo-electron Tomography

**DOI:** 10.1101/2024.01.05.574361

**Authors:** Lorenz Lamm, Ricardo D. Righetto, Tingying Peng

## Abstract

Cryo-electron tomography allows us to visualize and analyze the native cellular environment on a molecular level in 3D. To reliably study structures and interactions of proteins, they need to be accurately localized. Recent detection methods train a segmentation network and use post-processing to determine protein locations, often leading to inaccurate and inconsistent locations.

We present an end-to-end learning approach for more accurate protein center identification by introducing a differentiable, scoremap-guided Mean Shift clustering implementation. To make training computationally feasible, we sample random cluster points instead of processing the entire image.

We show that our Mean Shift loss leads to more accurate cluster center positions compared to the classical Dice loss. When combining these loss functions, we can enhance 3D protein shape preservation and improve clustering with more accurate, localization-focused score maps. In addition to improved protein localization, our method provides more efficient training with sparse ground truth annotations, due to our point sampling strategy.

## 1. INTRODUCTION

Cryo-electron tomography (Cryo-ET) is a promising imaging technique [1] that enables the study of cells in their native environment and in three dimensions. This innovative approach significantly advances our understanding of protein interactions in their native environment. A notable application is the determination of protein structures through subtomogram averaging (STA) [2], where small volumes are extracted around center positions of proteins within the tomogram, aligned, and then averaged to generate a high-resolution structure.

For STA, it is important to detect as many instances of the same protein as possible. These proteins must be precisely located to make STA efficient or even feasible. Therefore, determining initial center points as close as possible to the true protein centers is a critical step for the successful and efficient reconstruction of protein structures from native cells.

Classically, template matching [3] has been used for localizing proteins in Cryo-ET, and is still often used due to the lack of large public annotated datasets in Cryo-ET that could be used for training neural networks. The few available datasets often do not contain complete annotations and miss several true proteins [4]. Recently, template matching has been outperformed by new deep learning-based approaches in several cases [5, 6, 4, 7, 8]. Many of these methods [4, 7, 8] first train a neural network to segment protein shapes, and then use Mean Shift clustering [9] to extract protein center locations. Since the training is thus not focused on protein localization, resulting cluster centers may be inaccurate. Besides, these approaches require the often cumbersome generation of target maps depicting the protein shapes and do not directly utilize the protein center positions given by frameworks like template matching.

Mean shift clustering has been used for several deep learning tasks, including image segmentation [10, 11] and selfsupervised learning [12]. However, to our knowledge, it has not been proposed as a loss function for object center location, due to its non-differentiable nature.

We propose to integrate Mean Shift clustering into our network for end-to-end optimization of protein center locations. We introduce a score-weighted, differentiable Mean Shift module and attach it to a U-Net [13] architecture, enabling training with just protein center coordinates or combined with traditional segmentation loss. We show in multiple Cryo-ET datasets that this leads to more precise protein center locations, particularly in the case of non-spherical protein shapes. Furthermore, we show that our loss function yields good results even with incomplete ground truth annotations.

The code accompanying Mean Shift loss function and generating toy data can be accessed [here via GitHub](https://github.com/LorenzLamm/DeepFinder_MeanShift/tree/master).

## 2. METHODS

As shown in Figure 1A, existing approaches first train a UNet [13] to segment protein shapes, and then use Mean Shift clustering [9] as post-processing to extract protein center locations. We propose to train this workflow end-to-end by incorporating our differentiable variant of Mean Shift clustering into the architecture, enabling us to directly utilize ground truth (GT) protein center positions instead of (or in combination with) segmentation masks.

**Fig. 1.**
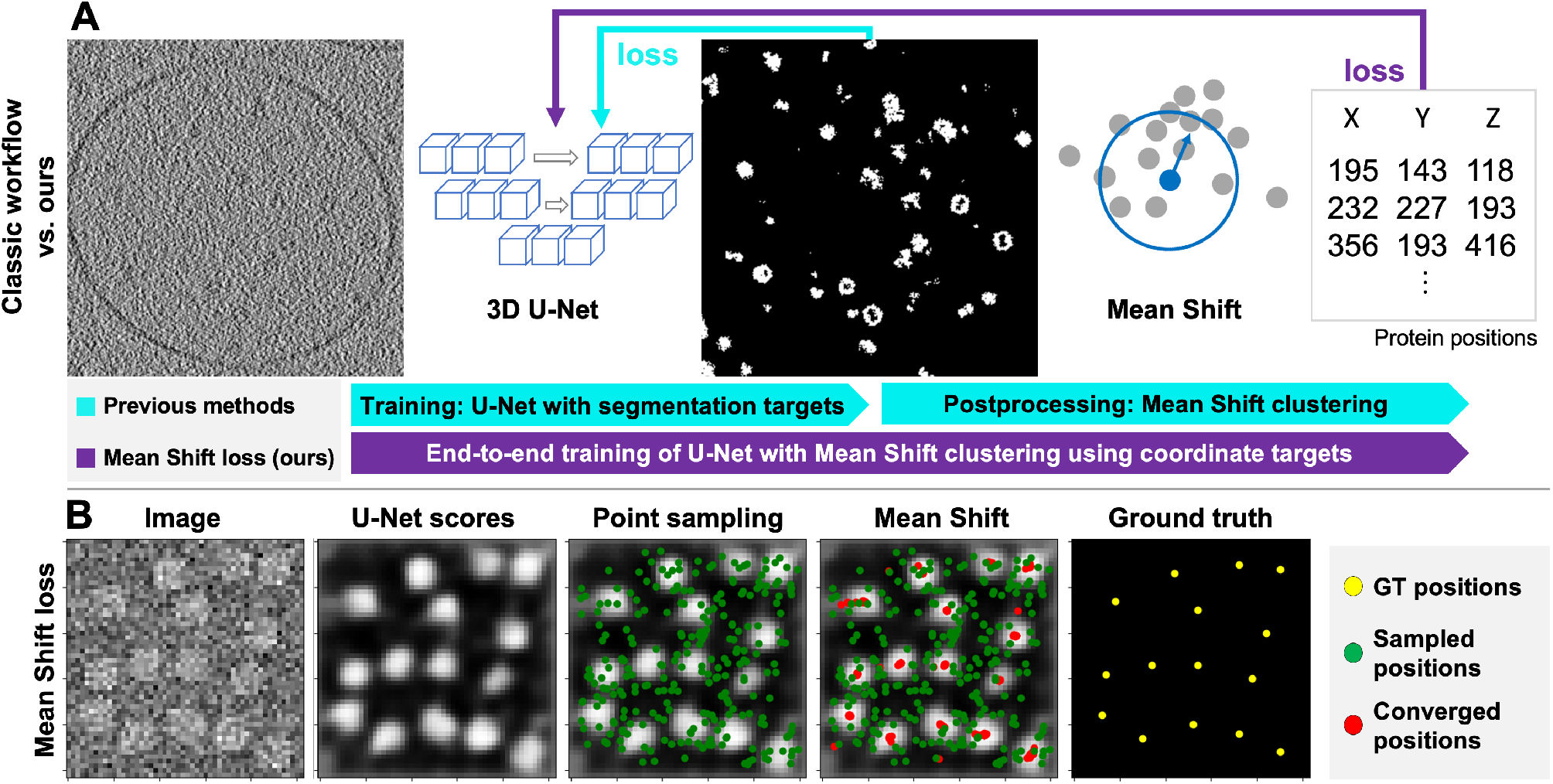
Mean Shift clustering as a loss function: **A:** Existing methods (e.g., [4, 8]) train a 3D U-Net to segment protein shapes. Subsequently, Mean Shift clustering gives protein center positions. Our Mean Shift loss allows to train this process end-to-end. **B:** Implementation of our Mean Shift loss: After computing U-Net score maps, random positions (green) are sampled around GT centers. Our differentiable Mean Shift clustering gives converged positions (red) from the sampled coordinates. These can be compared to GT positions (yellow) to compute a loss value.

### 2.1. Mean Shift clustering

Mean Shift clustering [9] is a clustering technique that iteratively shifts data points towards the densest part of a dataset. It is often used when the exact number of expected cluster centers is unknown, as its only adjustable parameter is the *bandwidth b*. The clustering processes each point separately by iteratively updating the point’s position by the weighted average of all points within radius *b* of the current point. All points thus converge to locally dense point regions.

### 2.2. Our differentiable Mean Shift clustering

Similar to [14], we implement Mean Shift using PyTorch on GPU in a batch-wise fashion. During inference, other methods [4, 8] perform thresholding of scoremaps to yield ini-tial cluster coordinates. Compared to that, during training, we randomly sample points within a certain radius around ground truth (GT) locations from the voxel grid (see Figure 1B). Then, we weight the sampled positions using the network-assigned scores of the corresponding voxels. Using these weights, we perform our score-guided Mean Shift clustering by iteratively computing the weighted averages of all points within bandwidth *b* of a sampled point *p*. For further efficiency, we limit the maximum of iterations to a low number (10 in all experiments). Algorithm 1 describes our score-weighted Mean Shift clustering in more detail.

#### Algorithm 1

Our differentiable Mean Shift loss

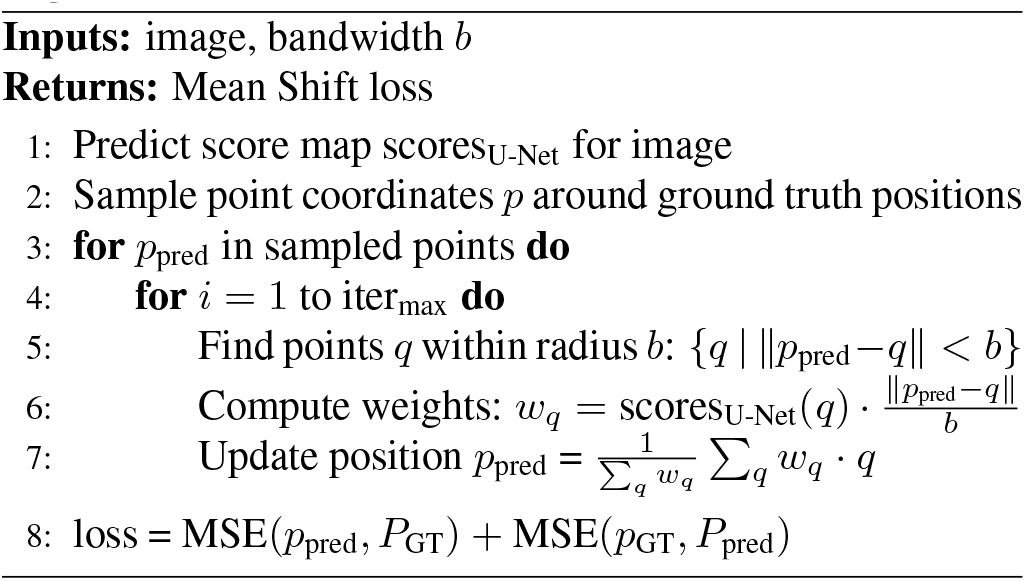

The introduction of network score-based weighting and random sampling of points without thresholding allows the backpropagation through the Mean Shift module and thus enables us to define our Mean Shift loss function. The advantages of sampling only a few positions (in practice, we use 256) around GT positions are twofold: First, it leads to a much more efficient clustering performance than processing all coordinates of the 3D patch. Second, this sampling allows us to focus our training on regions with available annotations: If a true protein position is not captured by the GT, we will not sample points close to this position and thus not severely distort the training process. Compared to that, classic segmentation metrics like Dice loss will be influenced strongly by to the false negative GT annotations.

After convergence, we have a set of predicted points *p*_pred_ ∈ *P*_pred_, and we use Mean Squared Error to compare to the GT positions *p*_GT_ ∈ *P*_GT_:

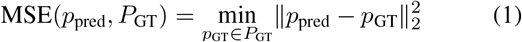

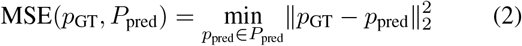

This ensures that GT positions are close to a predicted position, while prediction positions are close to a GT position.

### 2.3. Evaluation metrics

For evaluation, we define the average distances of predicted positions to their closest ground truth position and vice versa:

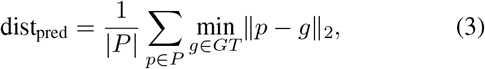

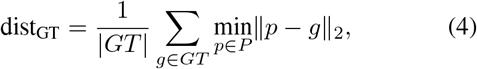

where *P* and *GT* are the sets of all predicted and GT positions, respectively. We also show the *F*_1_-score using different *hit*-radii: A GT position is counted as true positive (TP) with *hit*-radius 4 if a predicted position is within a radius of 4, and vice versa. Together with false positives (FP) and false negatives (FN), we compute precision, recall, and *F*_1_-score:

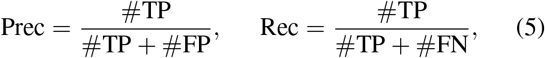

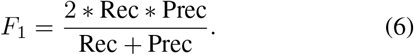

## 3. RESULTS

### 3.1. Datasets

We collected several datasets to benchmark our Mean Shift loss with the commonly used Dice loss. Figure 2 shows sample images of each dataset. As a proof of concept, we generated the *Sandclock* toy dataset by placing two spheres in opposite directions of a randomly drawn center point. Next, we used the synthetic *Shrec* Challenge Cryo-ET dataset [15], depicting proteins of different sizes in realistically simulated tomograms. Finally, we evaluated our approach using the *Ribo* dataset: a tomogram from an experimental dataset (EMPIAR10045 [16]) containing 3D locations of ribosomes.

**Fig. 2.**
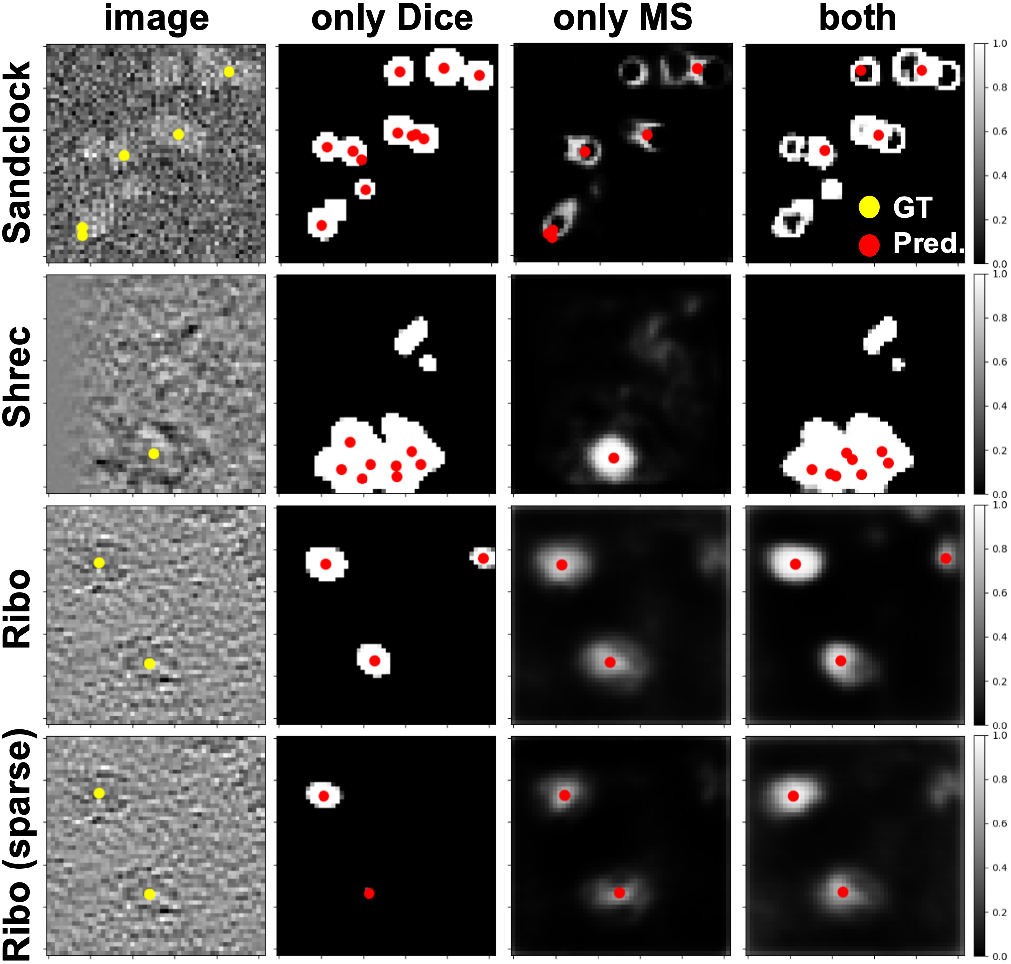
Images and predictions: 2D slices of the 3D patches of the *Sandclock, Shrec, Ribo*, and *Ribo (Sparse)* experiments. Shown are the raw input together with ground truth positions (yellow), as well as score maps for the experiments using only Dice as a loss functions, only our Mean Shift loss, or a combination. Predicted cluster centers are highlighted in red.

For the *Shrec* dataset, we sampled training (1558), validation (426) and test (440) patches that contained proteins from different tomograms. For the *Ribo* dataset, we generated nonoverlapping patches containing at least one protein, and split them into 50 training, 11 validation, and 11 test patches. For all datasets, we used a 3D patch size of 56^3^ both during training and evaluation.

### 3.2. Experimental evaluation

For each dataset, we performed training runs with Dice loss, Mean Shift loss, and their combination (Table 1, Figure 2), selecting the best model from 1000 epochs based on validation loss. We used a constant learning rate of 10^−5^ without weight decay or other regularization, and a bandwidth of 4 for Mean Shift clustering, tuned on the *Shrec* validation set.

**Table 1.** Results for different experiments. For each setting (*Sandclock, Shrec, Ribo, Ribo(sparse)*), we trained models using only Dice loss (_Dice_), only our Mean Shift loss (_MS_), and a combination of both (_both_). We show the means and standard deviations (5 training runs) of each predicted position’s distance to the closest ground truth position (dist_pred_), and vice versa (dist_GT_), as well as the F1-score with a *hit*-radius of 4.

| Experiment | $\text{dist}_{\text{pred}}$ | $\text{dist}_{\text{GT}}$ | $F1_{\text{rad4}}$ |
| --- | --- | --- | --- |
| Sandclock<sub>Dice</sub> | $4.02 \pm 0.01$ | $3.82 \pm 0.01$ | $0.46 \pm 0.01$ |
| Sandclock<sub>MS</sub> | **$2.43 \pm 0.17$** | $1.64 \pm 0.24$ | **$0.81 \pm 0.03$** |
| Sandclock<sub>both</sub> | $2.62 \pm 0.06$ | **$1.26 \pm 0.08$** | **$0.82 \pm 0.01$** |
| Shrec<sub>Dice</sub> | **$6.21 \pm 0.29$** | $2.09 \pm 0.16$ | $0.53 \pm 0.02$ |
| Shrec<sub>MS</sub> | $7.75 \pm 4.70$ | $3.99 \pm 1.70$ | **$0.70 \pm 0.10$** |
| Shrec<sub>both</sub> | $7.22 \pm 0.42$ | **$1.69 \pm 0.15$** | $0.55 \pm 0.01$ |
| Ribo<sub>Dice</sub> | $8.58 \pm 0.35$ | $4.03 \pm 0.04$ | $0.56 \pm 0.02$ |
| Ribo<sub>MS</sub> | $8.97 \pm 0.38$ | $4.82 \pm 0.85$ | $0.55 \pm 0.05$ |
| Ribo<sub>both</sub> | $8.44 \pm 0.23$ | **$3.54 \pm 0.10$** | **$0.61 \pm 0.02$** |
| Ribo(sparse)<sub>Dice</sub> | $8.24 \pm 0.68$ | $8.65 \pm 3.06$ | $0.45 \pm 0.10$ |
| Ribo(sparse)<sub>MS</sub> | $8.41 \pm 0.44$ | $6.46 \pm 1.54$ | **$0.53 \pm 0.05$** |
| Ribo(sparse)<sub>both</sub> | $8.87 \pm 0.49$ | **$4.13 \pm 0.47$** | **$0.55 \pm 0.05$** |

For the *Sandclock* dataset, we observe lower average distance values (dist_pred_), dist_GT_)) as well as better F1-scores when training with our Mean Shift loss or a combination. Figure 2 shows that while Dice loss offers more precise segmentations, it falls short in accurate cluster center identification. Conversely, our Mean Shift loss produces score maps that lead to more precise clustering and, consequently, more accurate protein center localization.

For the *Shrec* dataset, we observe mixed results: Dice loss or the combination show lower distance scores, but Mean Shift alone achieves the highest *F*_1_-score. The score maps from Dice loss training (Figure 2) more accurately predict protein shapes, but *Shrec*’s varying protein sizes lead to ambiguous cluster centers and potential protein oversampling, as uniform bandwidth clustering struggles with size variability. Conversely, Mean Shift loss generates score maps better suited for precise cluster center prediction, reflected in higher *F*_1_-scores. However, upon close inspection, we observed some significantly deviant outlier cluster centers, impacting distance values, as evident from the high standard deviations. The *F*_1_-score plot in Figure 3 further supports this, with Mean Shift loss achieving higher scores already at lower *hit*-thresholds, indicating overall accuracy despite outliers.

**Fig. 3.**
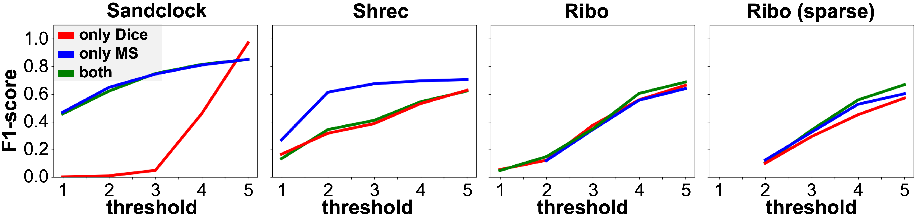
F1 scores for different *hit*-thresholds for all datasets and our three training settings using only Dice loss (red), only Mean Shift loss (blue), and a combination of both (green).

For the experimental dataset *Ribo*, we performed two experiments: First, we generated spherical masks around all ground truth positions and trained again using Dice loss, Mean Shift loss, and their combination. Here, we observe slightly improved distance scores and *F*_1_-scores when using the combined loss compared to only Dice loss. However, due to the roughly globular shape of the ribosomes, the advantage of using Mean Shift loss is not fully given.

Our second experiment *Ribo (sparse)* highlights our loss function’s ability to deal with sparse annotations: During training, we only used a single GT position and corresponding mask per patch to optimize our network. For the test set, we considered all GT positions again. While the performance using only Dice loss decreases notably (in particular dist_GT_, indicating many missed GT positions), training with the combined loss maintained similar results to full annotation training. This underscores our loss function’s capability to handle the sparse annotations common in Cryo-ET, where accurately localizing all proteins is often challenging.

## 4. CONCLUSION

To improve the accuracy of recent protein localization programs, we introduced a Mean Shift loss function that allows end-to-end training of a segmentation task with subsequent clustering. In order to use the originally non-differentiable Mean Shift clustering for training, we introduced a networkscore-based weighting to the clustering and implemented a point sampling scheme around GT positions to make the clustering computationally feasible. Using this Mean Shift loss, we can avoid tediously generating a segmentation target map and utilize ground truth locations directly.

We showed that, particularly for non-globular protein shapes, our loss function learns score maps that are more tailored towards a precise localization, compared to previous workflows with two separated steps to segment protein shapes and then perform independent clustering. Our point sampling strategy in the Mean Shift loss computation enhances robustness against sparsely annotated protein locations, a frequent issue in Cryo-ET.

In follow-up work, we would like to extend our loss function to a multi-class setting, and evaluate the benefits of the Mean Shift loss function on more experimental datasets (more diverse, non-globular protein shapes) and in more detail, e.g., by showing the effects of more accurate protein positions on downstream tasks like subtomogram averaging.

## 5. ACKNOWLEDGMENTS

L.L. acknowledges support from the Munich School for Data Science (MUDS) and a fellowship from the Boehringer Ingelheim Fonds. Calculations were performed at sciCORE (https://scicore.unibas.ch/) scientific computing center at the University of Basel. The authors acknowledge Ben Engel (University of Basel) for scientific discussions and access to computational resources.

## 6. COMPLIANCE WITH ETHICAL STANDARDS

This is a numerical simulation study for which no ethical approval was required.

